# MiR-212-3p functions as a tumor suppressor gene in group 3 medulloblastoma via targeting Nuclear Factor I/B (NFIB)

**DOI:** 10.1101/2021.04.15.440074

**Authors:** Naveenkumar Perumal, Ranjana K. Kanchan, David Doss, Noah Bastola, Pranita Atri, Ramakanth C. Venkata, Ishwor Thapa, Raghupathy Vengoji, Shailendra K. Maurya, David Klinkebiel, Geoffrey A. Talmon, Mohd W. Nasser, Surinder K. Batra, Sidharth Mahapatra

## Abstract

**Background:** Medulloblastoma (MB), the most frequent malignant pediatric brain tumor, is subdivided into four primary subgroups, wingless-type (WNT), sonic hedgehog (SHH), group 3, and group 4. Haploinsufficiency of chromosome 17p13.3 and c-Myc amplification distinguish high-risk group 3 tumors associated with rapid metastasis, recurrence and early mortality. We sought to identify the role of miR-212-3p, which resides on chromosome 17p13.3, in the pathophysiology of group 3 MB.

**Methods:** We first determined miR-212-3p expression in group 3 MB using several publicly-available datasets with confirmatory studies *in vitro*. We then identified epigenetic regulation by studying methylation and HDAC modifications along the promoter region. We used two systems for expression restoration, i.e. transient transfection or stable induction, to delineate miR-212-3p’s tumor suppressive and biochemical properties via assays assessing cancer proliferation, migration, invasion, colony formation, along with cell cycle and apoptosis analyses. We then compared MB and miR target databases to isolate a putative target whose biochemical and oncogenic properties were similarly elucidated using either transient silencing of target expression or stable induction of miR-212-3p.

**Results:** RNA expression analyses revealed dramatically reduced miR-212-3p levels in group 3 tumors and cell lines mainly through epigenetic silencing via histone modifications. Restoring miR-212-3p expression reduced *in vitro* cancer cell proliferation, migration, colony formation, and wound healing. Elevated miR-212-3p levels shifted c-Myc phosphorylation (from serine-62 to threonine-58), triggering destabilization and degradation; concurrently, its pro-apoptotic binding partners, i.e., Bin-1 and P19^ARF^, were upregulated with subsequent elevated apoptotic signals. Using a combination of transcriptomic data and dual luciferase assay, we isolated an oncogenic target of miR-212-3p, i.e. NFIB, a nuclear transcription factor implicated in metastasis and recurrence in various cancers. Increased expression of NFIB was confirmed in group 3 tumors, with poor survival shown in high-expressing patients. Transient NFIB silencing *in vitro* reduced cancer cell proliferation, colony formation, migration, and invasion. Concurrently, in group 3 MB cells, reduced medullosphere formation along with decreased expression of stem cell markers (Nanog, Oct4, Sox2, CD133) were noted.

**Conclusion:** These results substantiate the tumor-suppressive role of miR-212-3p in group 3 MB and provide a potential therapeutic oncogenic target implicated in metastasis and tumor recurrence.

## BACKGROUND

Medulloblastoma (MB), the most common malignant brain tumor of childhood, accounts for 20% of pediatric central nervous system (CNS) neoplasms with annual age-adjusted incidence ranging from 0.38 to 0.42 per 100,000 persons.^1,2^ Wingless-type (WNT), sonic hedgehog (SHH), group 3, and group 4 are the classic subgroups of MB with distinctive clinicopathologic and molecular features.^3^ The most aggressive tumors fall into group 3 (non-SHH/WNT MB), which account for approximately 25-30% of all MB cases and belong to a high-risk subgroup punctuated by haploinsufficiency of 17p (20-50% incidence), c-Myc amplification (15-20% incidence), metastases at diagnosis (30-40%), all resulting in very poor prognosis with <50% 5-year survivorship.^4–7^ Current treatment regimens involve surgical resection followed by a combination of craniospinal radiation and multi-agent chemotherapy, including vincristine, cisplatin, and either cyclophosphamide or lomustine.^8,9^ Recent studies have highlighted the importance of MB tumor-initiating (stem) cells, which are known to evade chemotherapeutic regimens, in tumorigenesis and recurrence.^10,11^ While the overall prognosis is poor in group 3 MB, recurrence can reduce 5-year survival to <10%.^5,12^ Thus, there is an urgent need to understand group 3 MB pathobiology and key signaling pathways involved in disease progression and tumor recurrence to provide a more accurate risk-adapted targeted treatment approach that can mitigate the dismal survivorship of these patients.

Frequent cytogenetic events target the 17p13.3 locus in group 3 MB and are associated with poor prognosis.^13–15^ Along this locus are multiple tumor suppressor genes and microRNAs (miRNAs), short nucleotide noncoding RNAs capable of inhibiting expression of target genes by preventing translation or promoting degradation.^16^ We recently reported on the tumor-suppressive properties of miR-1253, which lies on the terminal part of this locus, in MB.^17^ Similarly along this locus lies miR-212-3p, a tumor suppressor gene in various cancers, including lung cancer^18^, hepatocellular carcinoma^19^, prostate cancer^20^, and glioblastoma^21^. In colorectal cancer^22^ and nasopharyngeal carcinoma^23^, miR-212-3p targets MnSOD and Sox4, respectively, to prevent metastasis and invasion. In breast cancer, miR-212-3p regulates angiogenesis through Sp1 and VEGFA.^24^ It can also sensitize cetuximab-resistant cells to growth inhibition in Head and Neck Squamous Cell Carcinoma by targeting HBEGF.^25^ Other oncogenic targets of miR-212-3p include CTGF^19^, Engrailed-2^20^, and SGK3^21^, which are involved in cancer cell proliferation, migration, and invasion.

To date, no studies have examined the role of miR-212-3p in MB pathogenesis. Given its location on a highly afflicted chromosomal locus in high-risk MB, we hypothesized that miR-212-3p possesses tumor-suppressive properties. Here, we have focused on thoroughly elucidating the anti-neoplastic properties of miR-212-3p in group 3 MB and revealed a new oncogenic target, Nuclear Factor I/B (NFIB).

## METHODS

### Human tissue samples and molecular subgrouping of the MB tissues

Frozen and formalin-fixed paraffin-embedded (FFPE) samples of normal cerebellum (pediatric=12, adult=5) and pediatric MB specimens (WNT=1, SHH=7, grp 3=10, grp 4=14, unknown=7) were collected from the Children’s Hospital and Medical Center and the University of Nebraska Medical Center, Omaha. Tumor samples were sub-classified into four subgroups using genome-wide DNA methylome analysis (Illumina Methylation EPIC 850K bead arrays) as previously described.^17^ Normal cerebellum specimens were obtained at autopsy. For expression profile of HDACs (HDAC 1, 3, 5, 9), EZH2, and NFIB, we cross-analyzed the in-house MB dataset (Kanchan *et al.*, GSE148390) with publicly available MB datasets (Drusco *et al.*, GSE62381; and Weishaupt *et al*., GSE124814).^17,26,27^ For Kaplan-Meier Survival Analysis, we used the R2 database (Cavalli *et al.*, GSE85217).^4^

### Cell lines and cell culture

Human MB cell lines, D283 and D341, were purchased from ATCC; D425 was a kind gift from Darell Bigner (Duke University Medical Centre, Durham, NC); HDMB03 was a kind gift from Till Milde (Hopp Children’s Tumor Center, Heidelberg, Germany). DNA methylation profiling (New York University) and short tandem repeat (STR) DNA profiling (UNMC) were used to authenticate all cell lines. MB cell lines, i.e., D283, D341, D425, were maintained in DMEM medium with 10-20% FBS and 100 μg/mL penicillin/streptomycin; HDMB03 cells was cultured in RPMI with 10% FBS and 100 μg/mL penicillin/streptomycin. Untransformed Normal Human Astrocytes (NHA) and Human Neural progenitor cells (NPC) were purchased from Lonza Bioscience and maintained using respective basal medium supplemented with growth factors (Lonza Biosciences). Cell lines were maintained in 95% humidity, 37°C, 5% CO_2_.

### Reagents

MirVana™ miR-212-3p Mimic and scramble negative control were purchased from ThermoFisher Scientific. Silencing RNA for NFIB and EZH2 (Silencer™ Pre-Designed siRNA) and scrambled control (Silencer™ Negative Control) were obtained from ThermoFisher Scientific. Tet-On-miR-212-3p lentiviral vector and Tet3G (3rd generation) expression lentiviral vector were purchased from Vector Builder Inc.

### MicroRNA quantification

MiR-212-3p expression was quantified using TaqMan™ MicroRNA Reverse Transcription Kit and TaqMan™ Advanced miRNA Assay (Applied Biosystems). Isolated total RNA was reverse transcribed using a specific stem-loop primer for miR-212-3p and RNA U6B. After One-step TaqMan RT-PCR, miR-212-3p was quantified and normalized to RNU6B using the delta-delta Ct method.

### DNA methylation profiling

Genome-wide DNA methylome analysis was performed using the Illumina Methylation EPIC 850K bead arrays as previously described^28^. Genomic DNA was extracted from normal cerebellum and MB FFPE samples using RecoverAll™ Total Nucleic Acid Isolation Kit (Invitrogen). Results are presented as percent methylation at each CpG measured.

### De-methylation studies

HDMB03 cells (3×10^5^ in 6-well plates) were treated with global de-methylating agent 5-AzaC (5 μM) (5-Aza-2-deoxycytidine; Sigma). Following 96 hours of incubation, miR-212-3p expression was analyzed using TaqMan RT-PCR and presented as relative fold expression compared to control.

### Chromatin Immunoprecipitation Assay

The chromatin immunoprecipitation (ChIP) assay was performed with the Simplechip Enzymatic Chromatin IP kit (Cell Signaling Technology). Briefly, the digested cross-linked chromatin (10 μg) was subjected to immunoprecipitation with 5 μg of anti-H3K27me3 (Abcam), anti-H3K4me3 (Abcam), anti-H3K9me2 (Abcam), anti-H3K9Ac (Abcam), or mouse/rabbit IgG control (CST). Purified ChIP DNA was amplified using specific primers (Forward: 5′-GGAGTCCAGCTTCCTCTCCT-3′; and Reverse: 5′-GCTCCTGGGGGTCTTCAC-3’) detecting the CpG-enriched upstream promoter region of human miR-212-3p. Results are presented as relative enrichment normalized to respective input samples.

### MiR-212-3p expression restoration

Group 3 MB cells (3×10^5^) were seeded overnight and subsequently serum-starved for 4 hours prior to transient transfection using Lipofectamine 2000 (Invitrogen) for 6 hours in serum-free media with miR-212-3p mimic (25 nM and 50 nM) or scramble control (25 nM). Following incubation, fresh complete medium was added. For stable inducible system, HDMB03 cells were treated with or without Dox (500 ng/ml) in complete medium and incubated at 37°C, 5% CO_2_.

### Cell proliferation assay

MTT [3-(4, 5-dimethylthiazol-2-yl)-2, 5-diphenyl- 2H-tetrazolium bromide] and WST-1 [(4-[3-(4-Iodophenyl)-2-(4-nitro-phenyl)-2H-5-tetrazolio]-1,3-benzene sulfonate)] assays were used to determine cell viability in group 3 MB cells. After transfection/Dox treatment, assays were performed at 24-96 hours; absorbances were measured using a microplate reader at 570 nm (MTT assay) and 440 nm (WST-1 assay); data were analyzed using the SOFTMAX PRO software (Molecular Devices Corp.).

#### Colony formation assay

After transfection/Dox treatment, MB cells (1×10^3^ cells/well) were reseeded in 6-well plate and grown for 7-10 days in a humidified atmosphere (95% humidity) at 37°C and 5% CO_2_. Cells were washed, fixed with 2.5% methanol and stained with 0.5% crystal violet. Cell staining was dissolved using 10% acetic acid and quantified by measuring the absorbance at 590 nm.

### Cell migration and invasion assay

For trans-well migration/invasion assay, transfected/Dox treated stable cells (5×10^5^ cells) in serum-free media were seeded in the upper chamber of an insert (8 mm pore size; BD Bioscience) coated with Fibronectin (BD Bioscience) or Matrigel (Invasion Chamber Matrigel Matrix) followed by addition of a chemoattractant (10% FBS in complete media) to the lower chamber. After overnight incubation (16 hours), cells that migrated/invaded into the lower chamber were stained using Diff-Quik Stain Set (Siemens Healthcare Diagnostics, Inc.); images were captured using an EVOS FL Auto Imaging System (Life Technologies). Results were quantitated by taking an average cell count, measuring cell numbers from four-field/images/well (10× magnification).

### Wound healing assay

After transient transfection, cells (5×10^5^ cells/well) were plated in a 6-well plate, and upon reaching 80% confluence, a vertical scratch was made using a 10 μL pipette tip. For stable cells, 3×10^4^ cells/well were seeded in a culture-insert (ibidi culture-insert 2 well), and after overnight incubation culture-insert was removed and washed with PBS to remove non-adherent cells. Fresh growth medium (with and without Dox (500 ng/ml)) supplemented with 5% serum was added to the plates. The wound closure area was photographed at denoted time intervals using an EVOS FL Auto Imaging System (Life Technologies). Quantitative measurements (% wound closure) were determined by measuring three fields per well.

### Apoptosis and cell cycle Analysis

Annexin-V/Cy™5 (BD Biosciences) and propidium iodide (PI) (Roche Diagnostics) were used to measure apoptosis and cell cycle profile as previously described.^17^ Briefly, miR-212-3p mimic or scramble control transfected cells were incubated for 96 hours. Following incubation, cells were washed and resuspended in calcium-binding buffer consisting of Annexin-V/Cy™5 and PI (apoptosis assay) or fixed with 70% ethanol and stained with PI (cell cycle analysis). Stained cells were analyzed using a FACS Canto™ Flow cytometer (BD Bioscience).

### Western blotting

Protein lysates (30 μg/lane) were separated on 10% SDS-PAGE gels and transferred to PVDF membrane. Following blocking, target proteins were detected by probing overnight at 4°C with antibodies against: NFIB, Bin1 and p19^ARF^ (Abcam); PARP, cleaved PARP, cleaved caspase-3, caspase-3, cyclin D1, CDK4, CDK6, p-Akt-ser473, β-catenin, and CD133 (Cell Signaling Technologies); and total Akt, p-Erk and ERK (Santa Cruz Biotechnology). Then, membranes were washed and incubated with anti-mouse/rabbit IgG secondary antibody (Invitrogen) conjugated with horseradish peroxidase (HRP) at room temperature. Specific proteins were visualized using an enhanced chemiluminescence detection reagent (Pierce; ThermoFisher Scientific, Inc.).

### MiR-212 target prediction

MiRNA target prediction databases, i.e., Targetscan (http://www.targetscan.org) and mirDIP (http://ophid.utoronto.ca/mirDIP/) were used to determine the putative targets of miR-212-3p. Genes that were downregulated upon miR-212-3p restoration in HDMB03 (RNA sequencing analysis) were compared against them. Targets were chosen based on log_2_ fold change <−1 and *p* <0.05, yielding 37 common gene targets for miR-212-3p.

### Dual-luciferase reporter assay

Primers for 3’UTR-NFIB (Forward: 5’-GCTTGCTAGCTACACACCAGGGT-3’; and Reverse: 5’- TAGCAGTATAGGCTGGATA-3’) were designed using NCBI-Primer-BLAST tool (https://www.ncbi.nlm.nih.gov/tools/primer-blast/) and purchased from Eurofins. Following PCR amplification of NFIB (3’UTR), mutations were created within the seed sequence of NFIB. The resulting 3’UTR-Wild and 3’UTR-Mutant genes were inserted into the XbaI restriction site of pGL3-control vector (Promega) expressing firefly luciferase. Dual-luciferase reporter assay was performed on HDMB03 cells (3×10^5^ cells/well) seeded in 12-well plates. Following overnight incubation, cells were co-transfected with 3’UTR-wild-pGL3/3’UTR-Mutant-pGL3 plasmid, pRL-TK plasmid (Promega) expressing Renilla luciferase (internal control), and miR-212-3p mimic/scramble control for 48 hours. Luciferase activity was then measured using the Dual-Luciferase Reporter Assay System (Promega) with a Luminometer (Biotek). Results are presented as Luciferase activity (%), where firefly luciferase activity was normalized to Renilla luciferase activity (internal control) for each transfected sample.

### Immunohistochemistry

Unstained tissue slides (3 μm) were deparaffinized in xylene, rehydrated in a series of alcohol dilutions, and heated in citrate buffer (pH 6.2) for 20 minutes. Following antigen retrieval, slides were incubated with blocking buffer (Horse serum, Vector Labs) for 60 minutes at room temperature. Subsequently, anti-NFIB (1:100) primary antibody was added and incubated overnight at 4°C. After PBS wash, slides were incubated with secondary antibody (goat anti-rabbit/mouse conjugated with horseradish peroxidase) for 60 minutes at room temperature. Detection was performed with DAB Peroxidase Substrate Kit (Vector Labs) followed by counterstaining with hematoxylin. Sections were visualized under EVOS FL Auto Imaging System (Life Technologies). Then the tissues were scored according to the intensity of the dye color and the number of positive cells. The method for IHC score was as follows: 0, negative; 1, <25% positive tumor cells; 2, 25–50% positive tumor cells; 3, 50%-75% positive tumor cells; and 4, >75% positive tumor cells. Dye intensity was graded as 0 (no color), 1 (light yellow), 2 (light brown), or 3 (brown). Composite scores were derived from product of staining intensity and % positive cells (0–12).

### Sphere Formation Assay

For primary and secondary spheres, MB cells (4×10^5^ cells/well) were seeded in Ultra-Low Attachment 6-well plate (Corning™ Costar™ Microplates). Cells were maintained in stem cell media (DMEM/F12, 2% B27, 20□ ng/mL bFGF, 20 □ng/mL EGF, 40□U/ml penicillin, and 40□μg/ml streptomycin), and incubated for 10 days at 37°C, 5% CO_2_. Following incubation, primary spheres were dissociated into single cell suspension, reseeded, and allowed to grow for another 10 days to form secondary spheres. Spheres having diameter larger than 50 μm were counted.

### Statistical analysis

All experiments were conducted in triplicates. Values are presented as mean ± SD. Statistical analyses were performed using Prism 7.0b (GraphPad Software). For data normalization, control group was set at “1” and experimental groups were represented as fold-change compared to control with error bars reflecting deviation from mean triplicate measurements; statistical analyses were conducted prior to normalization. Differences between groups were compared using a two-tailed, unpaired Student’s t-test, Mann-Whitney U test or one-way analysis of variance followed by a least significant difference post hoc test. Statistical significance was established at \**p* <0.05, \*\**p* <0.01, and \*\*\**p* <0.001.

## RESULTS

### MiR-212-3p expression is epigenetically silenced by histone modification in group 3 MB

To explore the regulation of miR-212-3p in MB, we first conducted an *in silico* expression analysis of miR-212-3p in CSF samples of patients with MB (GSE62381, NC=14, MB=15), revealing reduced expression overall compared to normal cerebellum (**Figure 1A**). Subsequent expression examination in our local pediatric MB cohort (GSE148390) revealed specific downregulation in group 3 (n=9) and group 4 (n=14) MB tumors (**Figure 1A**). We validated these results using RT-PCR (**Figure 1B**), noting a near-abrogation of expression of miR-212-3p in group 3 MB samples (n=9) compared to pediatric cerebellum (n=10). Likewise, we revealed dysregulated expression of miR-212-3p in a panel of classic MB cell lines (group 3 MB – D341, D425, HDMB03; group 3/4 MB cell line – D283) compared to neural progenitor cells (NPC) using RT-PCR (**Figure 1C**). These results substantiate a decreased-to-absent expression of miR-212-3p in group 3 MB tumors and cell lines.

**Figure 1.**
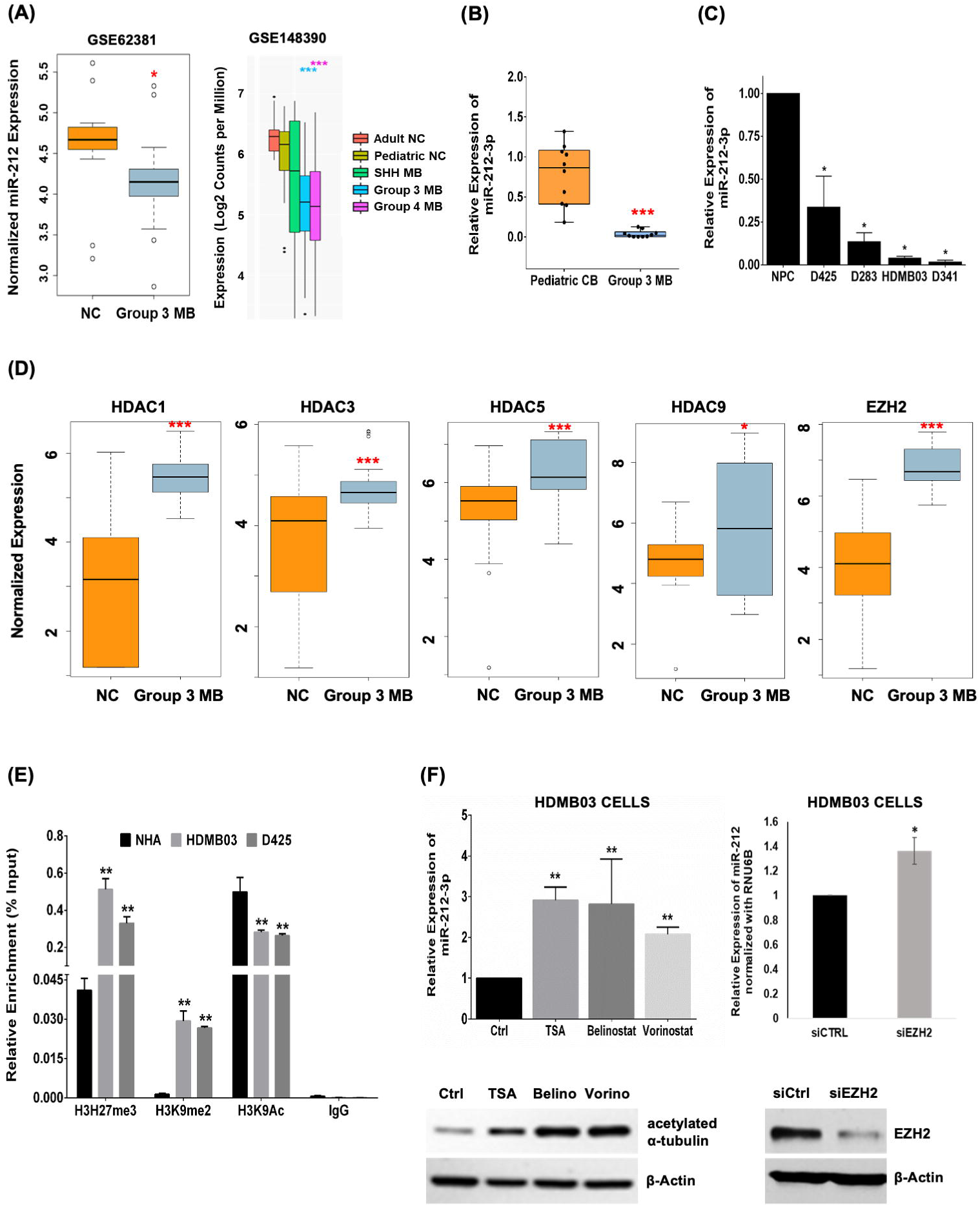
Expression and epigenetic regulation of miR-212-3p in group 3 MB tumors. **(A)** RNA sequencing analysis confirmed reduced expression of miR-212 in group 3 MB in two publicly-available MB datasets (Drusco *et al.*, GSE62381 and Kanchan *et al.*, GSE148390). **(B)** Results validated by RT-PCR normalized to U6b expression in group 3 MB tumor samples (n=10) compared to normal pediatric cerebellum (n=10). **(C)** RT-PCR analysis showing *in vitro* expression of miR-212-3p in classic MB cell lines (group 3: D341, D425, HDMB03; group 3/4: D283) compared to neural progenitor cells (NPC). RNU6B set as endogenous control. **(D)** Investigating histone modification as an epigenetic mechanism for miR-212 modulation in group 3 MB tumors (n=9) compared to pediatric cerebellum (n=10) via *in silico* expression of HDACs and EZH2 (Kanchan *et al.*, GSE148390). **(E)** ChIP-qRT-PCR analysis in group 3 MB cells demonstrates deregulation of histone modifications in the promoter region of miR-212-3p. ChIP grade histone mark antibodies to H3K27me3, H3K9me2 and H3K9Ac used; IgG antibody was negative control. **(F)** RT-PCR analysis showing miR-212-3p expression restoration after treatment with pan-HDAC inhibitors (TSA, 100 nM; Belinostat, 1 μM; and Vorinostat, 1 μM) and after inhibition using siRNA-EZH2 (20 nM) in HDMB03 cells. Western blotting analyses show increased acetylated α-tubulin levels in pan-HDAC inhibitors treated HDMB03 cells compared to vehicle control treated cells. β-actin served as an internal control. Data presented as mean ± SD from experiments done in triplicates and analyzed using Mann-Whitney U test (**A** and **D**) or Student’s t-test (**B, C, E** and **F**); \**p* <0.05, \*\**p* <0.01, \*\*\**p* <0.001.

To explore the mechanism for expression deregulation, we studied epigenetic silencing mechanisms, i.e. hypermethylation vs. histone modification. We initially performed DNA methylation profiling in our group 3 MB patient samples (n=6), but found a lack of perturbation to the methylation of the miR-212-3p promoter region when compared to normal pediatric cerebellum (n=4). *In vitro,* these findings were supported by a lack of expression restoration in 5-AzaC-treated (5 μM, 96 hour treatment) HDMB03 cells (**Supplementary Figure 1A**).

We then shifted our focus to histone modification-mediated epigenetic regulation. Our initial *in silico* analysis of HDAC expression revealed high expression of HDAC 1, 3, 5, 9, and EZH2 in both our local cohort (GSE148390, group 3 MB n=9) and in a larger group 3 MB cohort (GSE124814, group 3 MB n=211) when compared to normal cerebellum (n=10 and n=291, respectively) (**Figure 1D** and **Supplementary Figure 1B**). To confirm histone modification in the predicted miR-212-3p transcription start-site (TSS), histone mark patterns of group 3 MB cell lines (HDMB03 and D425) were compared with normal human astrocytes (NHA). CHIP-qRT-PCR analysis revealed significant differences in the methylation status of H3K27 and H3K9, and in the acetylation status of H3K9 in group 3 MB cell lines (**Figure 1E**). Specifically, HDMB03 and D425 cells, with baseline reduced miR-212-3p expression, showed enriched methylated H3K27 and H3K9, with a concomitant decrease in acetylated H3K9, compared to NHA. Consequently, treatment of the same cell lines with pan-HDAC inhibitors (TSA, 100 nM; Belinostat, 1 μM; Vorinostat,1 μM) substantially increased miR-212-3p expression compared to vehicle control (**Figure 1F** and **Supplementary Figure 1C**). Silencing EZH2 expression (siRNA-EZH2, 20 nM) accomplished the same (**Figure 1F**). Together, these findings strongly implicate histone modification (vs. hypermethylation) as a silencing mechanism for miR-212-3p in group 3 MB tumors.

### MiR-212-3p expression restoration inhibits group 3 MB cell line growth and proliferation

To highlight the tumor-suppressive properties of miR-212-3p in group 3 MB, we employed two methods to restore miR-212-3p expression, i.e. by transient transfection with miR-212-3p mimic or by Dox-inducible stable expression in group 3 MB cell lines. With successful miR-212-3p expression restoration (**Figure 2A**), cancer cell growth and proliferation were significantly impacted in a time- and dose-dependent manner (**Figure 2B**). Similarly, colony formation, transwell migration, and wound closure assays all recapitulated this anti-neoplastic phenotype with reduced colonies, cellular migration, and wound closure, respectively (**Figures 2C-E**). These data provide compelling *in vitro* evidence for the tumor-suppressive properties of miR-212-3p in group 3 MB.

**Figure 2.**
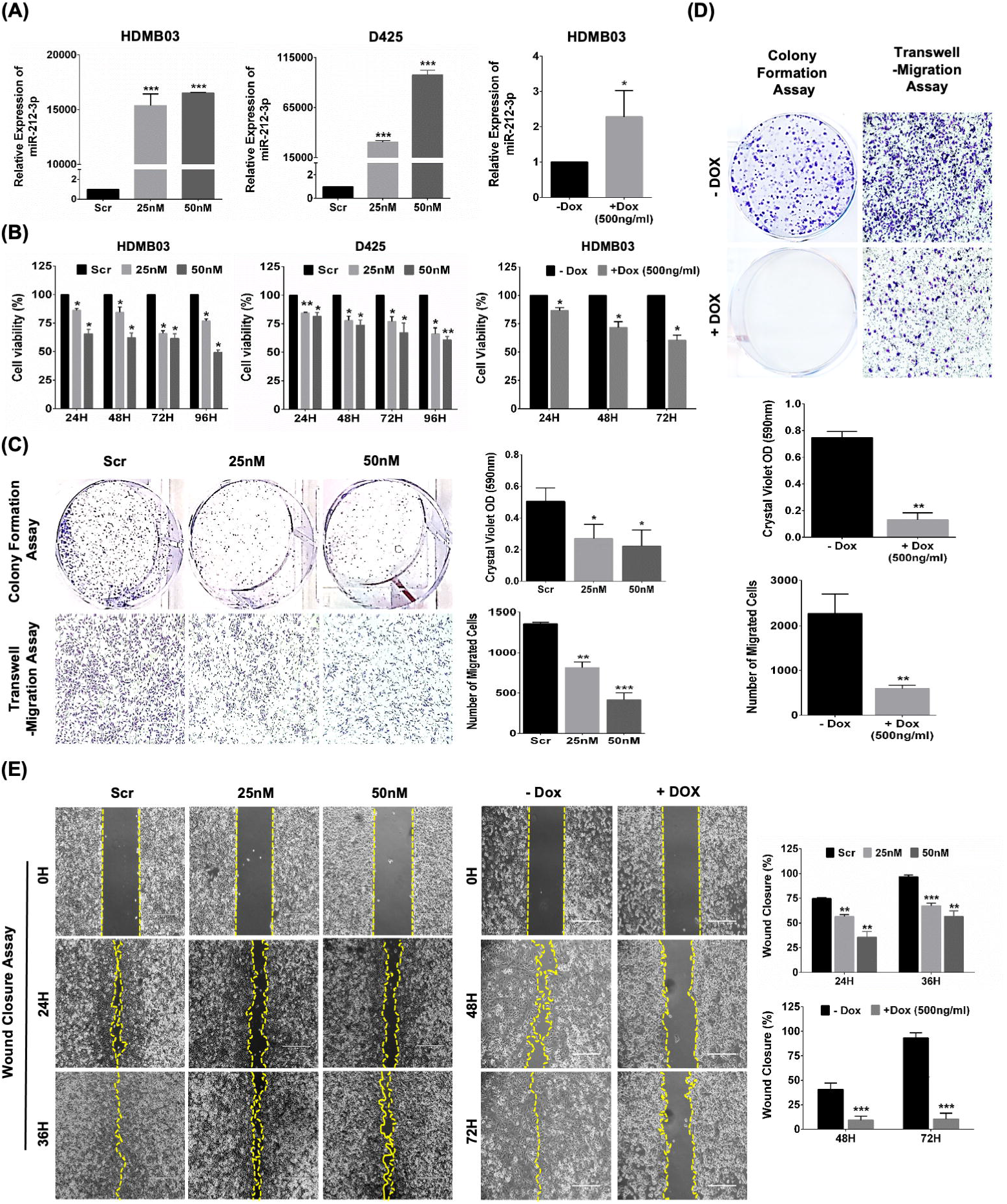
Effect of miR-212-3p expression on the neoplastic potential of group 3 MB cancer cells. **(A)** MiR-212-3p expression restoration via transient (HDMB03 and D425) and Dox inducible (HDMB03) systems was validated by RT-PCR. **(B)** Cell proliferation assays (MTT and WST-1) showed significantly reduced proliferative potential with both transient and stable expression of miR-212-3p in a time- and dose-dependent manner in group 3 MB cells. Similar growth-restrictive properties demonstrable via colony formation and transwell migration **(C and D)** and wound healing assays **(E)** with both transient and stable expression of miR-212-3p in HDMB03 cells. β-Actin served as an internal loading control. Data presented as mean ± SD from experiments done in triplicates and analyzed using Student’s t-test; \**p* <0.05, \*\**p* <0.01, \*\*\**p* <0.001. Scale bar: 100 μm.

### MiR-212-3p destabilizes c-Myc to favor cell cycle arrest and apoptosis in group 3 MB

Myc amplification is a cardinal high-risk feature of group 3 MB, and its phosphorylation status influences downstream tumor phenotype.^29,30^ More specifically, c-Myc phosphorylated at serine 62 increases stability leading to tumor aggressiveness, while phosphorylation at threonine 58 destabilizes the protein, leading to ubiquitin-mediated degradation and subsequent cellular apoptosis.^31,32^ In transiently transfected cells, miR-212-3p destabilized c-Myc by increasing phosphorylation of T58 and concurrently reducing phosphorylation at S62; however, in stably-expressing MB cells, total c-Myc levels were dramatically lowered (**Figure 3A**). Moreover, the active upstream kinases responsible for c-Myc phosphorylation at S62, i.e. p-Erk and p-Akt, were both significantly decreased in miR-212-3p expressing MB cells (**Figure 3A**). These data purport a c-Myc deregulatory function for miR-212-3p.

**Figure 3.**
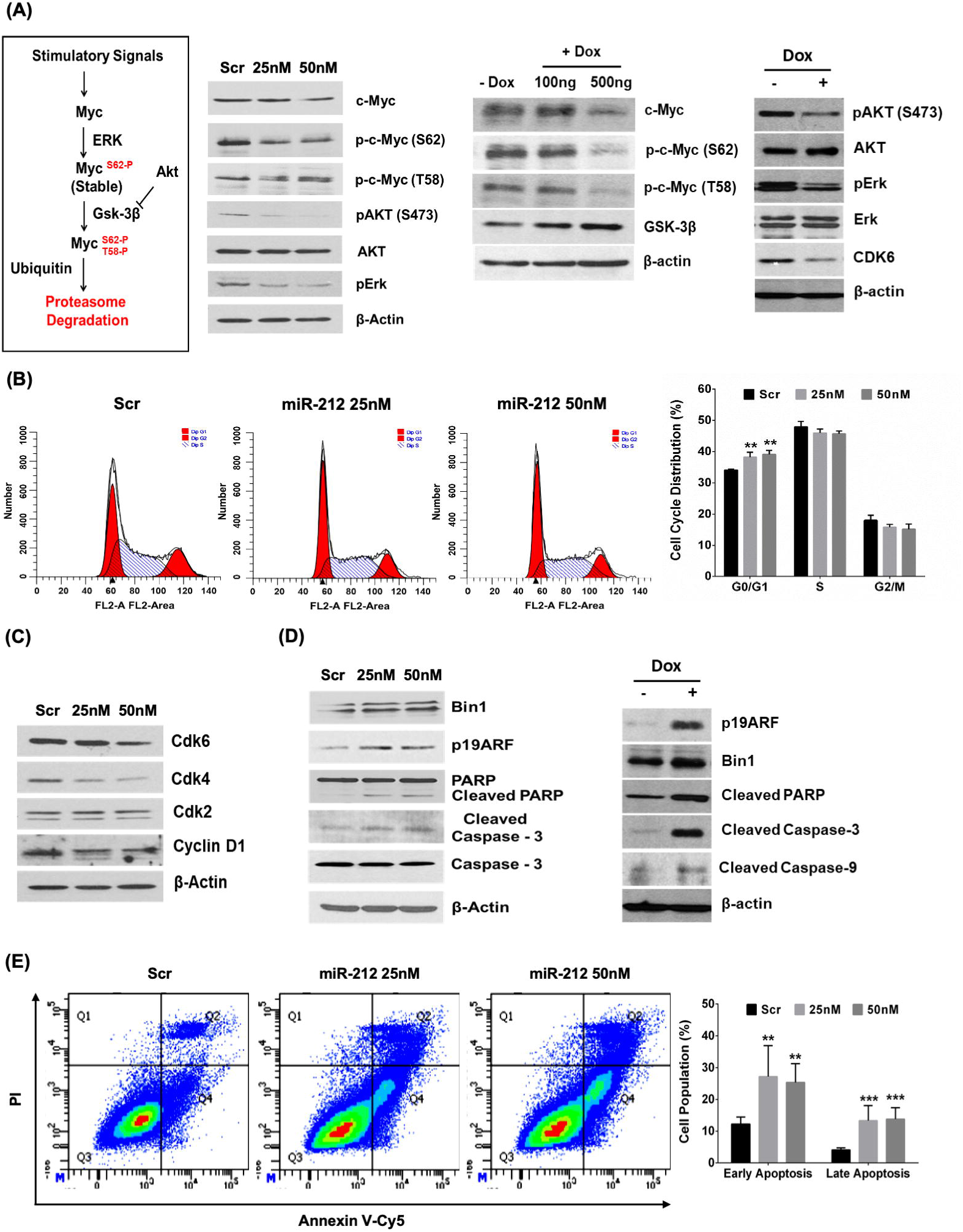
Effect of miR-212-3p expression on c-Myc regulation, cell cycle progression and apoptosis in group 3 MB cancer cells. **(A**) Western blotting analysis of c-Myc stimulatory signals showed a shift in c-Myc phosphorylation states from serine-62 (active form) to threonine-58 (inactive form) in miR-212-3p transiently transfected; while in Dox-treated stably-expressing HDMB03 cells, total c-Myc was reduced. Proliferation markers, i.e., p-Akt and p-Erk, upstream activators of c-Myc, were also significantly destabilized upon miR-212-3p restoration. **(B)** Cell cycle analysis by propidium iodide (PI) staining showed arrest at G_0_/G_1_ phase in miR-212-3p mimic transfected HDMB03 cells. **(C)** Supportive Western blotting analysis demonstrating reduced expression of G_0_/G_1_ regulatory checkpoint proteins, CDK4, CDK6, and cyclin D1, but not CDK2, in miR-212-3p mimic transfected HDMB03 cells. **(D)** Elevated expression of pro-apoptotic binding partners of c-Myc, i.e., Bin-1 and p19ARF, concurrent with apoptotic proteins (cleaved PARP and cleaved caspase-3) in miR-212-3p restored HDMB03 cells. **(E)** Annexin-Cy5 and PI staining confirmed increased apoptosis (early and late) in miR-212-3p mimic treated HDMB03 cells compared to scramble control. β-Actin served as an internal loading control. Data presented as mean ± SD from experiments done in triplicates and analyzed using Student’s t-test; \**p* <0.05, \*\**p* <0.01, \*\*\**p* <0.001.

We then focused on progression through the cell cycle, given the destabilization of c-Myc and a reduced proliferative phenotype noted in the presence of miR-212-3p. Not surprisingly, we revealed arrest in transiently-transfected miR-212-3p mimic-treated cells (25 nM and 50 nM) at the G_0_/G_1_ phase of the cell cycle in HDMB03 cells (**Figure 3B**). We confirmed the phase of arrest using Western blotting, which showed decreased expression of the complementary checkpoint markers, CDK4, CDK6, and cyclin D1 (**Figures 3A** and **3C**). In support, hierarchical clustering and pathways analysis revealed that miR-212-3p expression restoration led to enrichment of gene clusters (4, 5, 8) involved in regulation of cell cycle phase transition (**Supplementary Figure 2** and **Supplementary Table 2**).^33,34^

We concluded our study of the anti-neoplastic properties of miR-212-3p by analyzing effect on cancer cellular apoptosis. We first examined the expression of c-Myc binding partners that signal for apoptosis, i.e. Bin-1 and p19^ARF^, along with various pro-apoptotic proteins. MiR-212-3p restoration in HDMB03 cells significantly increased expression of Bin-1 and p19^ARF^ concurrent with rises in cleaved PARP, cleaved caspase 3 and 9 (**Figure 3D**). These results were validated using Annexin V-cy5/PI staining demonstrating a 2 and 3-fold increase in late and early apoptosis, respectively, compared to scrambled control (**Figure 3E**).

Taken together, our results identify an apoptotic mechanism for miR-212-3p, either by altering c-Myc phosphorylation states to destabilize c-Myc, by reducing total c-Myc expression, and/or by increasing the expression of its pro-apoptotic binding partners. In parallel, miR-212-3p plays a role in cell cycle arrest at the G_0_/G_1_ phase.

### NFIB is a downstream target of miR-212-3p

To identify oncogenic targets of miR-212-3p, we started with targets that are common to miRNA target prediction databases (TargetScan and mirDIP) and that are significantly downregulated (Log_2_ fold change <−1.0, *p* <0.05) by miR-212-3p expression restoration in HDMB03 cells (**Figure 4A**). This comparison revealed 37 putative targets whose expression and associated pathways were studied in two group 3 MB patient cohorts (**Figure 4B** and **Supplementary Figure 3**). Out of the 37 common targets, fourteen were upregulated in group 3 MB (summarized in **Supplementary Table 1**). NFIB was chosen for further study based on the following characteristics that bestow a high likelihood of oncogenic potential in group 3 MB: (i) There are two conserved binding sites for miR-212-3p on the 3′UTR of NFIB mRNA (TargetScan); (ii) expression of NFIB was significantly increased in group 3 tumors (GSE148390; n=9, and GSE124814; n=211) compared to normal cerebellum (n=10, and n=291, respectively) (**Figure 4C**); (iii) Kaplan-Meier Survival Analysis showed poor survival in high-expressing patients (Cavalli *et al.*, GSE85217, via R2 database)^4^ (**Figure 4D**); (iv) immunohistochemical staining of NFIB in group 3 MB tissues (n=9) revealed intense nuclear staining when compared to normal cerebellum (n=8) (**Figure 4E**); and (v) high expression was shown in a panel of group 3 MB cell lines (HDMB03, D425 and D341) compared to NHA (**Figure 4F**). We further validated this target by the dual-luciferase assay (**Figure 4G**) and showed significant attrition in miR-212-3p mimic treated HDMB03 cells at the transcription (**Figures 4H**) and translation level (**Figure 4I**). In this manner, we identified NFIB as an oncogenic target of miR-212-3p.

**Figure 4.**
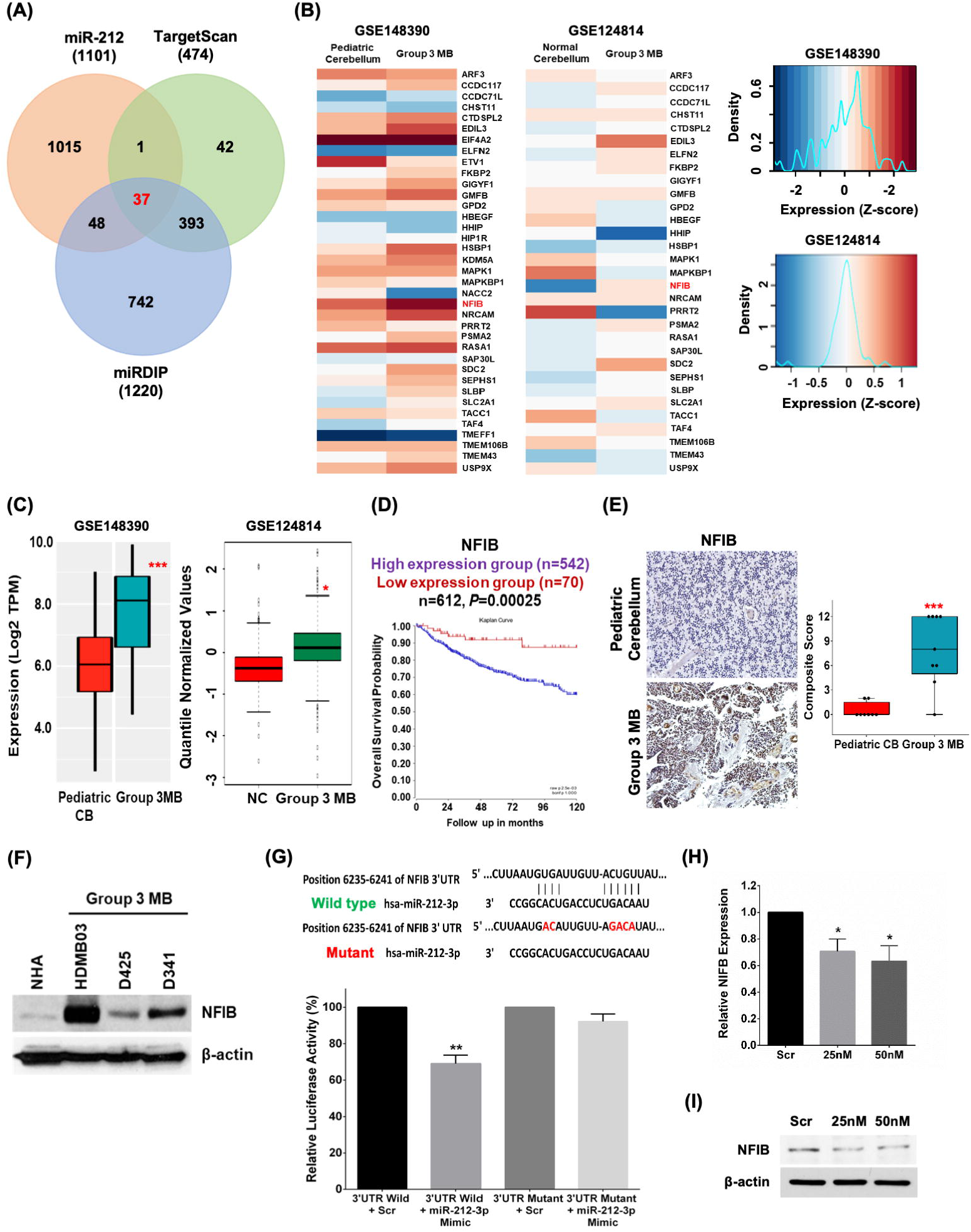
Isolating oncogenic targets of miR-212-3p in group 3 MB tumors. **(A)** Venn diagram showing 37 common targets between miRNA target prediction databases (Targetscan and miRDip) and genes downregulated (Log_2_ fold change <−1.0, *p* <0.05) in miR-212 expressing HDMB03 cells (via RNA sequencing analysis). **(B)** Expression heat map of common target in group 3 MB data sets (GSE148390, n=9, and GSE124814, n=211) compared to normal cerebellum (n=10 and n=291, respectively). NFIB was chosen for further study as a strong putative oncogenic target of miR-212-3p based on: **(C)** elevated expression of NFIB in group 3 MB data sets (GSE148390 and GSE124814); **(D)** Kaplan-Meier Survival Analysis (R2 database, Cavalli *et al.*, GSE85217) demonstrating poor survival in high-expressing NFIB MB patients (n=542) compared to low-expressing patients (n=70); **(E)** immunohistochemical staining of NFIB showing intense nuclear expression in group 3 MB patient samples (n=9) when compared to normal cerebellum (n=8); and **(F)** Western blotting showing significantly increased expression of NFIB in classic group 3 MB cell lines (HDMB03, D425, and D341). **(G)** Dual-luciferase assay confirmed NFIB as a direct target of miR-212-3p. **(H)** RT-PCR and Western blotting analysis showed significantly decreased expression of NFIB in miR-212-3p transiently transfected HDMB03 cells. β-actin served as an internal loading control. Data presented as mean ± SD from experiments done in triplicates and analyzed using Mann-Whitney U test **(C)** or Student’s t-test (**E, G,** and **H**); \**p* <0.05, \*\**p* <0.01, \*\*\**p* <0.001. Scale bar: 100 μm.

### NFIB possesses oncogenic potential in group 3 MB cancer cells

Nuclear Factor I/B (NFIB) belongs to the nuclear factor I (NFI) family of transcription and replication proteins that recognize palindromic sequences on various promoters capable of activating transcription and replication throughout organ development.^35,36^ An oncogenic role for NFIB has been shown in triple-negative breast cancer (TNBC), small cell lung cancer (SCLC), colorectal and gastric cancers, and melanoma by enhancing tumor growth, epithelial-mesenchymal transition (EMT), migration and invasion.^37–41^

With this prior evidence, we elucidated the oncogenic role of NFIB in group 3 MB cancer cells by studying the effect of silencing expression on proliferation, transwell-migration and invasion. Successful silencing of NFIB (siRNA-NFIB, 20 nM) in HDMB03 cells for 48 hours (**Figure 5C** and **5D**) resulted in a significant reduction in cell proliferation in a time-dependent manner (**Figure 5A**). Trans-well migration and invasion assays demonstrated similar reductions in cell migration and invasion (**Figure 5B**).

**Fig. 5.**
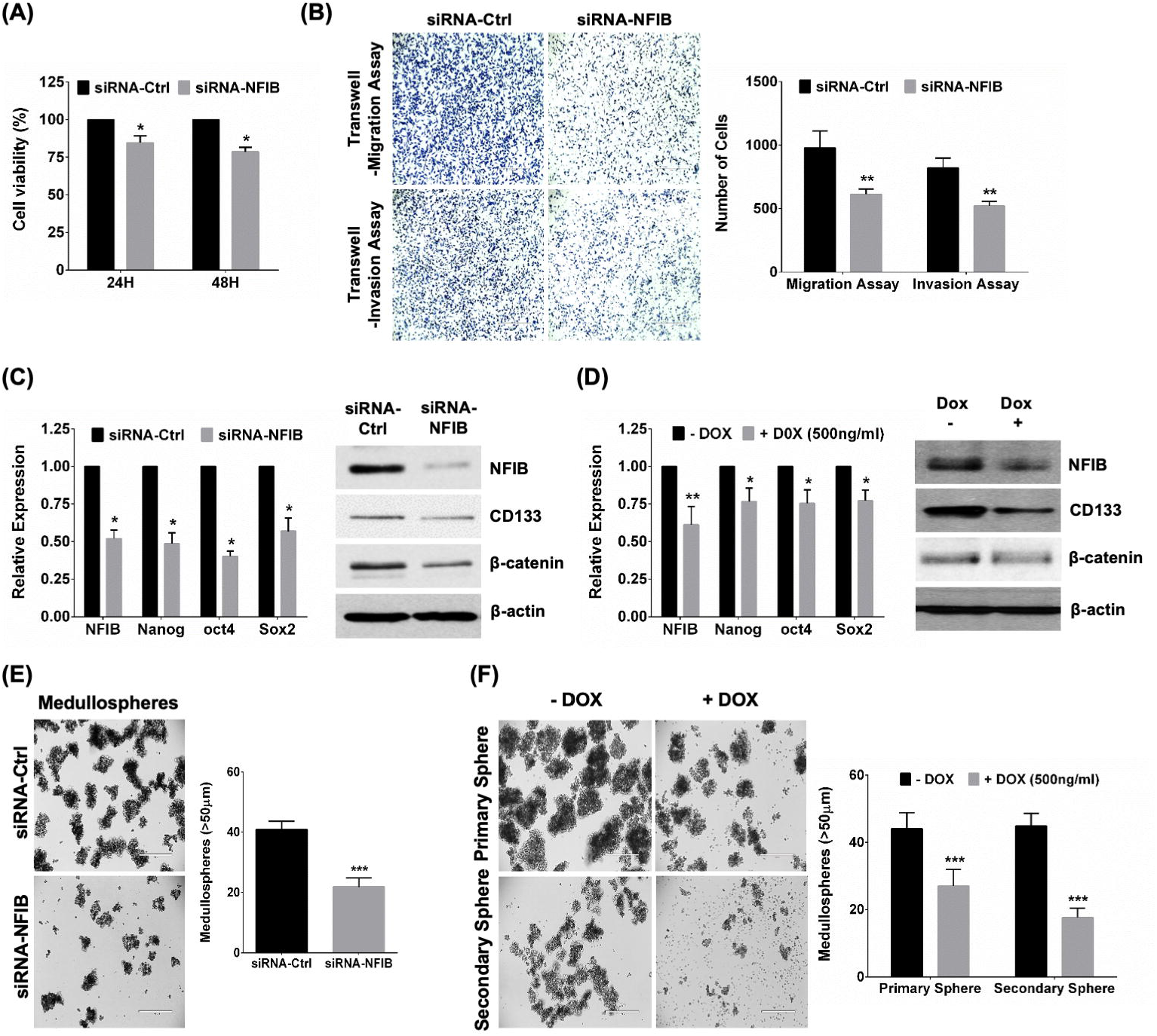
Effect of NFIB silencing on cancer cell phenotype in group 3 MB. **(A)** Transient silencing of NFIB significantly decreased cell viability in siRNA-NFIB (20 nM) treated HDMB03 cells compared to siRNA-ctrl cells. **(B)** Trans-well migration/invasion assay also showed decreased cell number in siRNA-NFIB treated cells compared to siRNA-ctrl. RT-PCR and Western blotting demonstrated reduced stem cell markers, i.e. Nanog, Oct4, Sox2, CD133 and β-catenin, in **(C)** siRNA-NFIB treated HDMB03 cells and **(D)** Dox (500 ng/ml) treated stably-expressing miR-212-3p HDMB03 cells. **(E** and **F)** Medullosphere assays demonstrated dramatically reduced sphere formation in siRNA-NFIB-treated and in Dox-treated stably-expressing miR-212-3p HDMB03 cells, respectively. β-Actin served as an internal loading control. Data presented as mean ± SD from experiments done in triplicates and analyzed by Student’s t-test; \**p* <0.05; \*\**p* <0.01; \*\*\**p* <0.001. Scale bar: 100 μm.

Mechanistically, NFIB overexpression was shown to increase chromatin accessibility, promote expression of pro-metastatic genes, and drive metastasis in SCLC tumors.^39,42,43^ Previous studies in SCLC revealed NFIB as a downstream target of c-Myc, which directly regulated its expression and contributed to rapid metastases.^44^ Given the direct inhibitory effect of miR-212-3p on c-Myc and NFIB’s promoting role in metastasis, we sought to study the effect of the miR-212-3p/NFIB axis on cancer stem cell (CSC) maintenance and self-renewal. We first analyzed the expression of stem cell markers in miR-212-3p stably-expressing and NFIB-silenced HDMB03 cells. RT-PCR and Western blotting analyses revealed concurrent deregulation in the expression of several stem cell markers, including CD133, β-catenin, Sox-2, Oct-4, and Nanog (**Figures 5C** and **5D**). CSCs self-renewal ability was further assessed through tumor sphere-forming assay. Again, in both NFIB-silenced and miR-212-3p stably-expressing cancer cells, substantially decreased numbers of spheres were noted (**Figures 5E** and **5F**). These results not only substantiate miR-212-3p’s role in CSC maintenance but also implicate NFIB as a strong oncogenic target of miR-212-3p in group 3 MB.

## DISCUSSION

Haploinsufficiency of chromosome 17p bestows a high-risk feature that distinguishes the non-SHH/WNT groups 3 and 4 MB tumors.^4,5,7^ Several loci populated by tumor suppressor genes can be found within the afflicted short arm. Studies have linked few genes residing on the terminal locus (17p13.3) with MB, such as ROX/MNT^14^ or HIC1^45^, but none have studied micro RNAs on this locus. We previously elucidated the anti-neoplastic properties and oncogenic targets of miR-1253, a micro RNA found on the terminal part of this locus, in MB.^17^ In the present study, we have described the tumor-suppressive properties of miR-212-3p in group 3 MB; we have additionally identified a target with strong oncogenic potential.

Initial *in silico* analyses of publicly-available group 3 MB datasets confirmed a significantly deregulated expression profile for miR-212-3p compared to normal cerebellum. These findings were recapitulated both in pediatric tumor samples (*ex vivo*) and in classic group 3 cell lines (*in vitro*). In many cancers, microRNAs undergo epigenetic silencing via hypermethylation along CpG islands or chromatin rearrangement through histone modifications.^17,46^ Methylation profiling in high-risk tumors showed no differences in methylation pattern of the miR-212-3p locus between tumors and normal tissue. Contrarily, histone modifications at critical lysine residues within miR-212-3p provided compelling evidence for an epigenetic silencing mechanism. More specifically, increased methylation at H3K27 and H3K9 and a concomitant decline in acetylation of H3K9 was shown by ChIP-RT-PCR. Either treatment with HDAC inhibitors or EZH2 silencing reliably restored miR-212-3p expression. Together, these data not only revealed miR-212-3p silencing in group 3 tumors, but also assigned an epigenetic mechanism via histone modifications. Our findings align with prior reported patterns of miR-212-3p epigenetic silencing in lung cancer.^18^

Restoring miR-212-3p expression in group 3 MB cells by either transient transfections or stable induction resulted in a cadre of anti-neoplastic effects. First, cancer cell proliferation, wound healing, migration, and colony formation were significantly reduced. Subsequently, cell proliferative markers, p-Akt and p-Erk, were downregulated in miR-212-3p restored cells, eventually leading to cell cycle arrest (G_0_/G_1_ cell cycle phase) with dysregulated expression of checkpoint regulatory kinases, CDK4, CDK6, and cyclin D1. Next, we revealed deregulation of c-Myc in miR-212-3p restored cells, with a shift in phosphorylation patterns from S62 (active, favoring proliferation) to T58 (inactive, favoring apoptosis), and a decrease in total c-Myc levels. Of note, p-Akt and p-Erk are upstream kinases that phosphorylate c-Myc at S62, rendering it resistant to degradation^31,32^; as we have shown, miR-212-3p restricts this process. Moreover, in miR-212-3p-expressing cells, key apoptotic binding partners of c-Myc, p19^ARF^ and BIN1, were elevated in parallel with cleaved PARP, cleaved caspase□3, and cleaved caspase-9, resulting in a high apoptotic signal in cancer cells. Similar results were observed in our previous investigation of miR-1253 on this locus.^17^ Given the high association between c-Myc amplification and poor prognosis in group 3 MB^4,7^, our data suggest a plausible mechanism contributing to high-risk MB aggressiveness, wherein silencing miR-212-3p can lift its regulation of c-Myc, allowing for unchecked Myc-driven signaling. As prior, our results align with studies on miR-212-3p in colorectal cancer^22^, nasopharyngeal carcinoma^23^, and lung cancer^47^ where similar effects were seen on cancer cell proliferation, migration and invasion, cell cycle arrest, and apoptosis.

We next identified an oncogenic target of miR-212-3p using *in silico*, *ex vivo,* and *in vitro* approaches. We, thus, revealed Nuclear Factor I/B (NFIB), a transcription factor that can bind to specific overlapping DNA repeat sequences (5’-TTGGCNNNNNGCCAA-3’) throughout the genome to activate transcription and replication, most notably in normal lung and brain development.^35,36^ In fact, the 14 target genes of miR-212 isolated via high-throughput analyses of independent MB datasets seemed most intricately involved in the same (**Supplementary Figure 3B**). Notwithstanding its high expression in group 3 tumors associated with poor survival, we showed that silencing NFIB expression dramatically decreased cancer cell proliferation, migration, and invasion *in vitro*. Similar effects have been shown for NFIB in gastric cancers^37^, TNBC^38^, SCLC^39^, and colorectal cancer^40^. In colorectal cancer, melanoma, and gastric cancers, overexpression of NFIB was associated with epithelial-mesenchymal transition (EMT), migration and invasion.^37,40,41^ In melanoma, specifically, NFIB-targeted upregulation of EZH2 led to the epigenetic silencing of MITF, promoting a highly invasive phenotype.^41^ In SCLC, NFIB overexpression cooperated with Rb/p53 deletion to increase chromatin accessibility to pro-metastatic genes. Moreover, c-Myc was shown to regulate NFIB in SCLC and further contribute to rapid metastases.^44^

Metastases at diagnosis and tumor recurrence are cardinal prognostic features contributing to dramatically increased mortality in group 3 MB.^4–7,29,30^ MB recurrent tumors, either in the primary or metastatic site, derive from a subpopulation of cancer stem cells (MBSCs), which can evade current chemotherapeutics and radiation therapy.^48^ MBSCs express stemness markers, such as CD133, CD15, and Sox2, and can possess an inordinate capability to form aggressive tumors with increased self-renewal ability, facilitating MB relapse and rapid demise.^49,50^ In NFIB-silenced and miR-212-3p restored group 3 MB cells, we noted decreased expression of stemness markers, i.e. CD133, Sox2, Oct4, Nanog, and β-catenin, concurrent with reduced tumor cell self-renewal capacity as evidenced by sphere-forming assays. These results introduce miR-212-3p’s role in hampering MBSC maintenance and self-renewal, possibly through NFIB regulation.

Intriguingly, we have shown EZH2 as an upstream regulator of miR-212-3p; we have also revealed miR-212-3p-mediated de-regulation of c-Myc. At the core of these critical contributors to tumor aggressiveness seems to be NFIB. Whether NFIB plays a decisive role in triggering neoplastic transformation in group 3 MB by upregulating EZH2 in group 3 tumors, thereby silencing miR-212-3p which in turn lifts c-Myc regulation, is an important topic of further study.

## CONCLUSIONS

This study has uncovered a novel tumor suppressor gene in group 3 MB, i.e. miR-212-3p. We have shown expression silencing by histone modifications, as opposed to hypermethylation. The anti-neoplastic properties of miR-212-3p are augmented by destabilization of c-Myc, cancer cellular arrest at G_0_/G_1_ cycle, and robust apoptosis. By targeting NFIB, a well-studied metastatic driver^42^, a regulator of EZH2^41^, and downstream effector of c-Myc^44^, miR-212-3p decreases cancer cell aggressiveness, stem cell maintenance, and renewal. Using miRNA-mediated therapeutics may provide the ideal approach to not only addressing group 3 MB tumor aggressiveness but also unburdening young patients from the harmful side□effects of current cytotoxic therapies.

## Abbreviations

3’UTR: 3’ untranslated region
5-AzaC: 5-Aza-2′-deoxycytidine
AKT: Protein kinase B
ANOVA: analysis of variance
Bin-1: Myc box-dependent-interacting protein 1
c-Myc: Avian myelocytomatosis virus oncogene cellular homolog
CB: cerebellum
CDK4: Cyclin-dependent kinase 4
CDK6: Cyclin-dependent kinase 6
CNS: central nervous system
MBSC: medulloblastoma stem like cells
CTGF: Connective tissue growth factor
DNA: deoxyribonucleic acid
EMT: epithelial-mesenchymal transition
EZH2: Enhancer of zeste homolog 2
FACS: fluorescence-activated cell sorting
FFPE: formalin-fixed paraffin-embedded
FISH: fluorescence *in situ* hybridization
HIC1: hypermethylated in cancer-1
HCC: Hepatocellular carcinoma
HDAC: histone de-acetylase
HRP: horseradish peroxidase
i(17q): isochromosome 17q
IHC: immunohistochemistry
IRB: Institutional Review Board
MB: medulloblastoma
miR: microRNA
miR-1253: microRNA 1253
miR-212-3p: microRNA 212-3p
MITF: Microphthalmia-associated transcription factor
MNT: Max network transcription factor
MTT: 3-(4, 5-dimethylthiazol-2-yl)-2, 5-diphenyl- 2H-tetrazolium bromide
NFIB: Nuclear factor I/B
Nl: normal
Non-SHH/WNT: Non-Sonic Hedgehog/Non-Wingless
NSCLC: Non-small cell lung cancer
Oct4: Octamer-binding transcription factor-4
p19^ARF^: p19 alternative reading frame protein
p-Akt: Phosphorylated protein kinase B
PARP: Poly ADP ribose polymerase
PCR: polymerase chain reaction
Ped: pediatric
p-Erk: Phosphorylated extracellular signal related kinase
PI3K: Phosphoinositide 3-kinase
RNA: ribonucleic acid
SCLC: non-small cell lung cancer
SHH: Sonic Hedgehog
SD: standard deviation
SGK3: Serum and glucocorticoid-inducible kinase 3
SMYD4: SET And MYND domain containing 4
Sox-2: Sry-box transcription factor-2
TCF7L2: Transcription factor 7-like 2
TNBC: triple-negative breast cancer
TSA: Tricostatin A
WNT: Wingless

## DECLARATIONS

### Ethics approval and consent to participate

Studies had prior approval from the Institutional Review Board of the University of Nebraska Medical Center (IRB # 561-16-EP). Due to exempt status, informed consent was waived.

### Consent for publication

Not applicable

### Availability of data and materials

The datasets used during the current study (GSE148390, GSE62381, GSE124814, and GSE85217) are available in the Gene Expression Omnibus (GEO). Data generated in the current study are available from the corresponding author on reasonable request.

### Authors’ contributions

NP, MWN, SKB, and SM designed the study. NP and RKK performed the experiments. DD, PA, RCV, and IT provided bioinformatics support for the study. NB and NP contributed to the isolation of putative targets of miR-212-3p. DK performed and analyzed epigenetic silencing studies. RV and SKM assisted in *in vivo* studies. GT provided support as the Pathologist on the study. NP, RKK, DD, PA, RCV, IT, MWN, SKB, and SM contributed significantly to the interpretation of the data. NP wrote the first draft of the manuscript. DD, RV, SKB and SM revised the manuscript. All authors read and approved the final manuscript.

## Acknowledgements

We would like to thank Dr. Amar Singh’s Lab for the provision of supplies to conduct luciferase assay. We would like to acknowledge Drs. Deborah Perry and Michael Punsoni for assisting with procuring tumor samples for the study. We would further like to acknowledge the contribution of our funding sources to this study.

---

**Supplementary Figure 1: Exploring epigenetic silencing of miR-212-3p in group 3 MB tumors. (A)** DNA methylation profile of group 3 (n=6) and 4 (n=11) MB patient tumors showing lack of perturbations to methylation in the promoter region of miR-212-3p compared to normal pediatric cerebellum (n=4); further demonstrable *in vitro* by a lack of expression restoration with de-methylation by 5-AzaC (5 μM) in HDMB03 cells. **(B)** Recapitulation of elevated HDAC and EZH2 expression complementing prior *in silico* data in a larger group 3 MB cohort (Weishaupt H *et al.*, GSE124814; n=211) compared to normal cerebellum (n=291). **(C)** RT-PCR analysis in pan-HDAC-treated (TSA, 10 nM; Belinostat, 1 μM; and Vorinostat, 1 μM) D425 cells showing elevated expression of miR-212-3p. RNU6B set as an endogenous control. Western blotting analysis showed increased acetylated α-tubulin in pan-HDAC inhibitors treated D425 cells. β-actin served as an internal control. Results presented as mean ± SD from experiments done in triplicates and analyzed using Student’s t-test **(A and C)** or Mann-Whiteny U test **(B)**; \**p* <0.05, \*\**p* <0.01, \*\*\**p* <0.001.

**Supplementary Figure 2: Deregulated pathways associated with miR-212-3p silencing in group MB. (A)** Bubble plot (FunSet plot) representing 12 clusters of significantly enriched GO biological processes. FunSet employs hypergeometric test to perform the enrichments and additionally uses semantic similarity measure with Aggregate Information Content (AIC) index to cluster highly similar GO terms.^34^ **(B)** Enriched biological pathways associated with miR-212-3p silencing identified using Enrichr (https://maayanlab.cloud/Enrichr/).^33^

**Supplementary Figure 3: Putative oncogenic targets of miR-212-3p and their associated pathways in group 3 MB tumors. (A)** Expression analysis by RNA Sequencing of isolated miR-212-3p targets (37) in group 3 MB (GSE148390, n=9) compared to normal pediatric cerebellum (n=10). Of those observed to be significantly elevated in medulloblastoma were the following 14 genes: *CCDC71L*, *CTDSPL2*, *EDIL3*, *GIGYF1*, *HSBP1*, *KDM5A*, *NFIB*, *NRCAM*, *PSMA2*, *SDC2*, *SEPHS1*, *TAF4*, *TMEM43*, and *USP9X*. **(B)** Enriched cellular/biological pathways associated with these targets identified using with FunSet tool.^34^ Box plots represent gene expression (Log_2_ Transcripts Per Million) and analyzed using Mann-Whiteny U test; \**p* <0.05, \*\**p* <0.01, \*\*\**p* <0.001. Pathways represented as an inverted bar graph based on the −log_10_ FDR values with threshold *p* <0.05 (red dashed line).

**Supplementary Table 1: Isolating a strong oncogenic target of miR-212-3p.** Further analysis of 14 putative miR-212-3p targets by examining conserved binding sites, confirming elevated expression in 2 independent MB cohorts, and examining effect of high expression on survival spotlighted NFIB as a pivotal oncogenic target in group 3 MB. In the table, ‘x’ denotes inclusion criteria, and ‘-’ denotes no expression data found in dataset.

**Supplementary Table 2: Genes (and associated pathways) enriched by miR-212-3p expression restoration in HDMB03 cells.** Detailed gene list for the 12 clusters of significantly enriched GO biological processes represented in **Supplementary Figure 2A**. Clusters were selected based on −log_10_ FDR values with threshold *p* <0.05.

