## Supplementary figures and images for "MiR-212-3p functions as a tumor suppressor gene in group 3 medulloblastoma via targeting Nuclear Factor I/B (NFIB)"

### Supplementary Figure 1

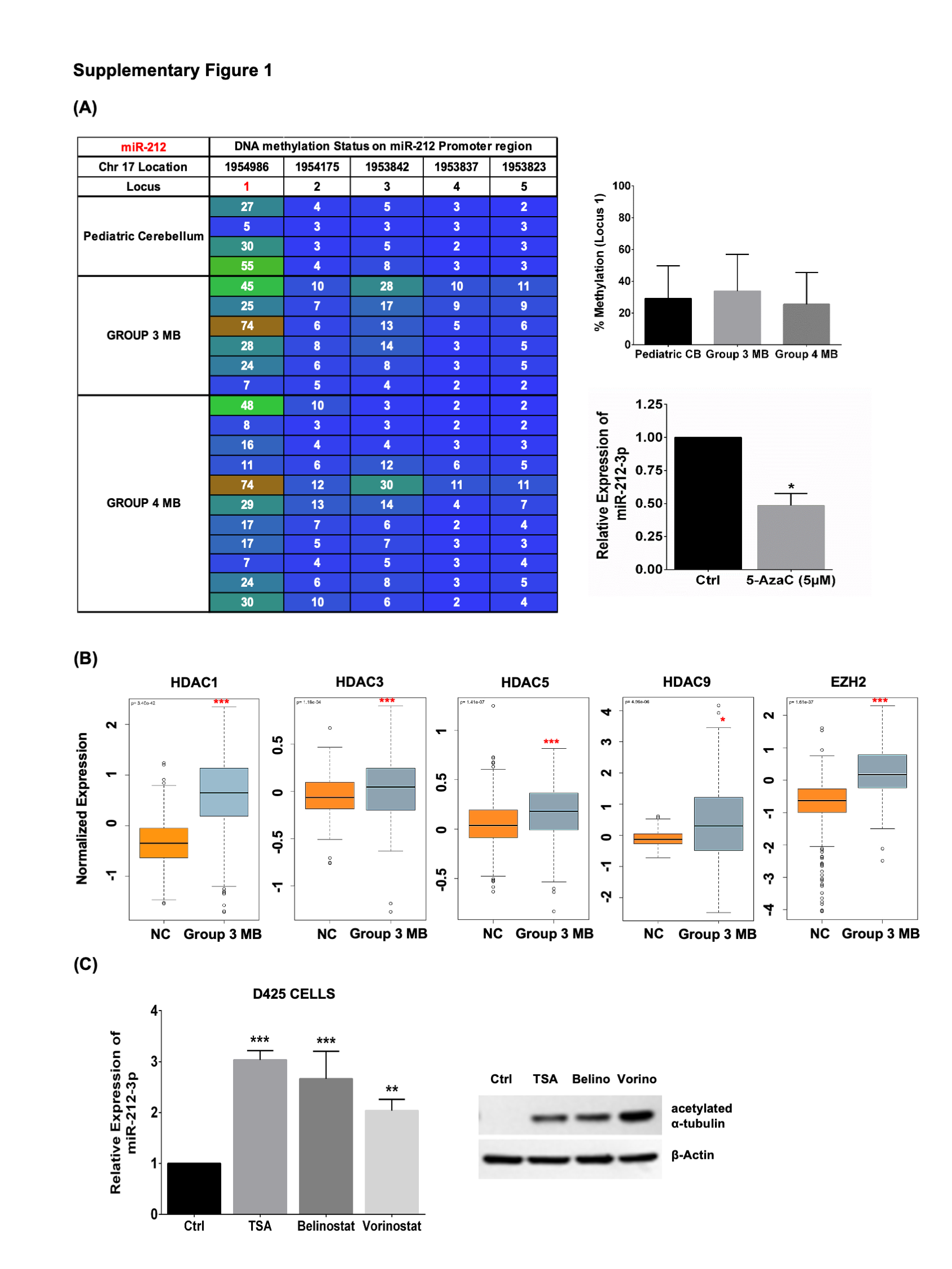

### Supplementary Figure 2

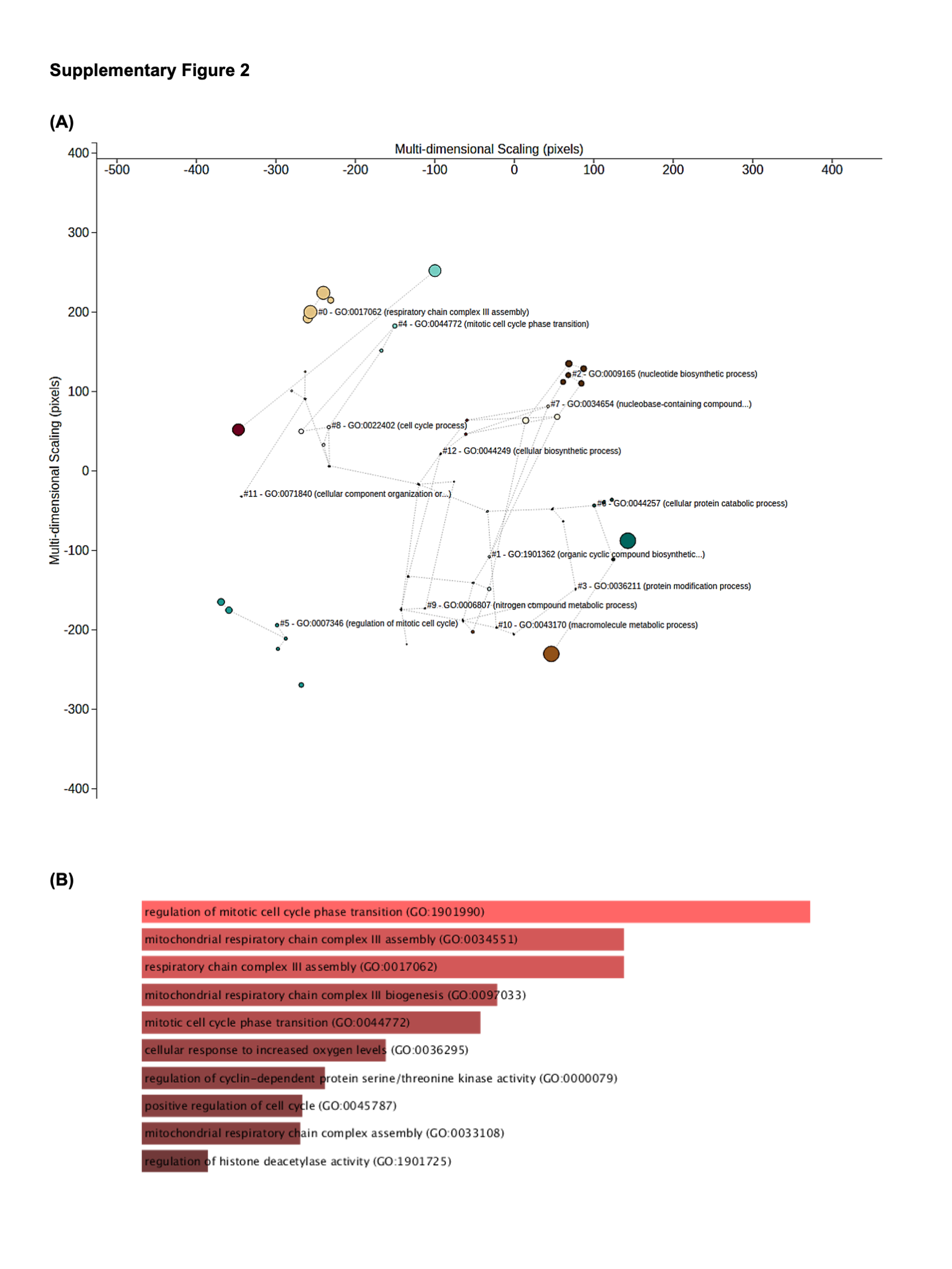

### Supplementary Figure 3

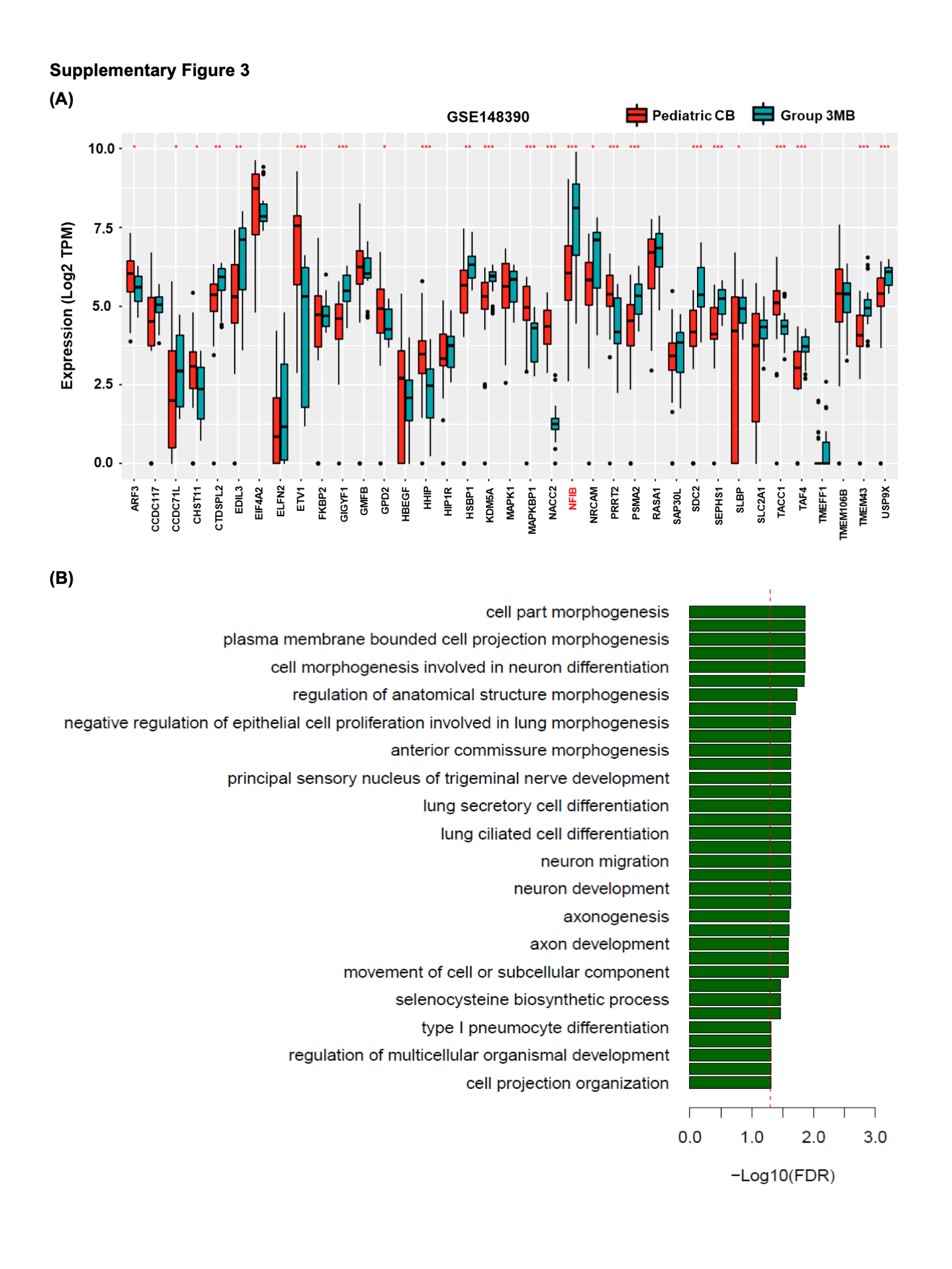

### Supplementary Table 1

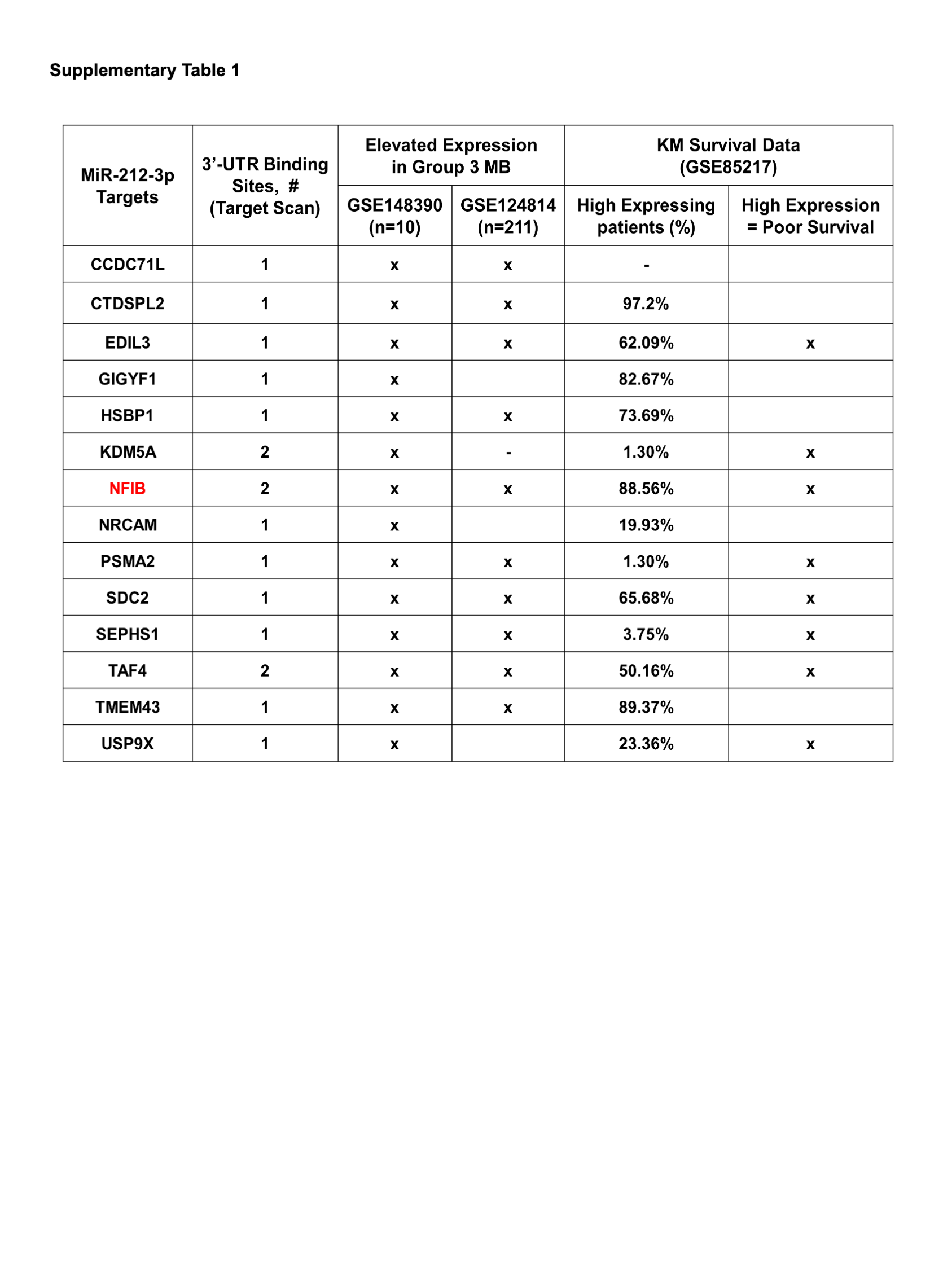
